# Porphyromonas gingivalis fuels colorectal cancer through CHI3L1-mediated iNKT cell-driven immune evasion

**DOI:** 10.1101/2023.12.29.573607

**Authors:** Angélica Díaz-Basabe, Georgia Lattanzi, Federica Perillo, Chiara Amoroso, Alberto Baeri, Andrea Farini, Yvan Torrente, Giuseppe Penna, Maria Rescigno, Michele Ghidini, Elisa Cassinotti, Ludovica Baldari, Luigi Boni, Maurizio Vecchi, Flavio Caprioli, Federica Facciotti, Francesco Strati

**Affiliations:** Department of Experimental Oncology, European Institute of Oncology IRCCS, Milan, Italy; Department of Oncology and Hemato-oncology, Università degli Studi di Milano, Milan, Italy; Gastroenterology and Endoscopy Unit, Fondazione IRCCS Cà Granda, Ospedale Maggiore Policlinico, Milan, Italy; Department of Biotechnology and Biosciences, University of Milano-Bicocca, Milan, Italy; Neurology Unit, Fondazione IRCCS Ca’ Granda Ospedale Maggiore Policlinico, Milan, Italy; Centro Dino Ferrari, Department of Pathophysiology and Transplantation, Università degli Studi di Milano, Milano, Italy; IRCCS Humanitas Research Hospital, Rozzano, Milan, Italy; Department of Biomedical Sciences, Humanitas University, Milan, Italy; Medical Oncology, Fondazione IRCCS Ca’ Granda, Ospedale Maggiore Policlinico, Milan, Italy; Department of General and Minimally Invasive Surgery, Fondazione IRCCS Ca’ Granda, Ospedale Maggiore Policlinico, Milan, Italy; Department of Pathophysiology and Transplantation, Università degli Studi di Milano, Milan, Italy

**Author notes:** Correspondence to: Francesco Strati, and Federica Facciotti,; Piazza della Scienza 2, Department of Biotechnology and Biosciences, University of Milano-Bicocca, 20126 Milano, Italy. These authors contributed equally (Angélica Díaz-Basabe, Georgia Lattanzi). These authors contributed equally (Federica Facciotti, Francesco Strati).

**Keywords:** iNKT cells, CRC, *Porphyromonas gingivalis*, CHI3L1

## Abstract

The interaction between the gut microbiota and invariant Natural Killer T (iNKT) cells plays a pivotal role in colorectal cancer (CRC). *Porphyromonas gingivalis* is a keystone oral pathogen associated with CRC. The oral pathobiont *Fusobacterium nucleatum* influences the anti-tumour functions of CRC-infiltrating iNKT cells. However, the impact of other oral bacteria, like *P. gingivalis*, on their activation status remains unexplored. In this study, we demonstrate that mucosa-associated *P. gingivalis* induces a protumour phenotype in iNKT cells, subsequently influencing the composition of mononuclear-phagocyte cells within the tumour microenvironment in CRC. Mechanistically, *in vivo* and *in vitro* experiments show that *P. gingivalis* reduces the cytotoxic functions of iNKT cells, hampering the iNKT cell lytic machinery though increased expression of chitinase 3-like-1 protein (CHI3L1). Neutralization of CHI3L1 effectively restores iNKT cell cytotoxic functions suggesting a therapeutic potential to reactivate iNKT cell-mediated antitumour immunity. In conclusion, our data demonstrate how *P. gingivalis* accelerates CRC progression by inducing iNKT cells to upregulate CHI3L1, thus impairing iNKT cell cytotoxicity and promoting host tumour immune evasion.

## Introduction

Colorectal cancer (CRC) is the third most prevalent cancer worldwide and the second leading cause of cancer-related death (*1*). The mutational landscape and the mechanisms of tumour initiation in CRC have been widely described, but colon carcinogenesis also depends on the interaction between cancer cells and the tumour microenvironment (TME) (*2*). Indeed, the polarization and activation profiles of immune cells within the TME is highly informative to predict CRC patient survival or their response to therapy, highlighting the importance of the inflammatory microenvironment for CRC tumorigenesis (*2*). Microbiota-elicited inflammation is an important contributor to CRC pathogenesis regardless of pre-cancer inflammatory history (*3*). Pro-carcinogenic bacteria are able to initiate and promote colon cancer, partly through mechanisms that are not fully understood (*4*). *Porphyromonas gingivalis* is an opportunistic oral pathogen associated with different inflammatory diseases and cancers (*5, 6*) and specifically enriched in CRC patients (*7, 8*). *P. gingivalis* accelerates epithelial cell proliferation through the MAPK/ERK signalling pathway (*9*) and upregulates the expression of senescence and proinflammatory genes through the local production of butyrate (*10*). Moreover, *P. gingivalis* promotes CRC immune subversion through activation of the hematopoietic NOD-like receptor protein 3 inflammasome in tumour-infiltrating myeloid cells (*11*). Recently, we demonstrated that tumour-infiltrating invariant Natural Killer T (iNKT) cells favour a proinflammatory yet immunosuppressive TME in CRC by sensing *Fusobacterium nucleatum*, an opportunistic pathogen of the oral cavity (*12*). Indeed, iNKT cell conditioning by *F. nucleatum* induces the iNKT cell-mediated recruitment of immunosuppressive PMN-MDSCs-like neutrophils, supporting CRC progression (*12*). iNKT cells are lipid-specific, evolutionary conserved, T lymphocytes (*13*) that participate in cancer immune surveillance, including CRC (*14*). iNKT cell functional impairment in the TME is associated with poor overall survival in solid and hematologic tumours but the environmental factors modulating their functionality are incompletely elucidated (*14*). Here, we address the contribution of *P. gingivalis* to the activation status and functions of tumour-infiltrating iNKT cells in CRC. We demonstrate that *P. gingivalis* imprint a protumour phenotype to iNKT cells which in turn affect the intratumour mononuclear-phagocyte cell landscape. Mechanistically, *P. gingivalis* impairs iNKT cell cytotoxicity by preventing the correct formation of the iNKT cell lytic machinery though the upregulation of chitinase 3-like-1 protein (CHI3L1) while promoting recruitment of neutrophils within the TME.

## Results

### Intestinal colonization by *Porphyromonas gingivalis* promotes CRC through iNKT cells

We have recently demonstrated that tumour-infiltrating iNKT cells contribute to CRC tumorigenesis through interactions with *F. nucleatum* (*12*). Given the common pathogenic features of *P. gingivalis* (*Pg*) and *F. nucleatum* (*5, 15*), we hypothesized that *Pg* might similarly induce a protumour phenotype in iNKT cells. Thus, we classified a cohort of CRC patients based on the presence of *Pg* in their mucosal-associated microbiota, which had been characterized in our prior study (*12*). We observed a significant enrichment of tumour-infiltrating iNKT cells and neutrophils in *Pg*^positive^ as compared to *Pg*^negative^ CRC patients (Figure 1A and 1B). The enrichment of both neutrophils and iNKT cells is consistent with our previous observations which underscore the iNKT cell-mediated recruitment of tumour-associated neutrophils (TANs) in CRC lesions (*12*).

**Figure 1:**
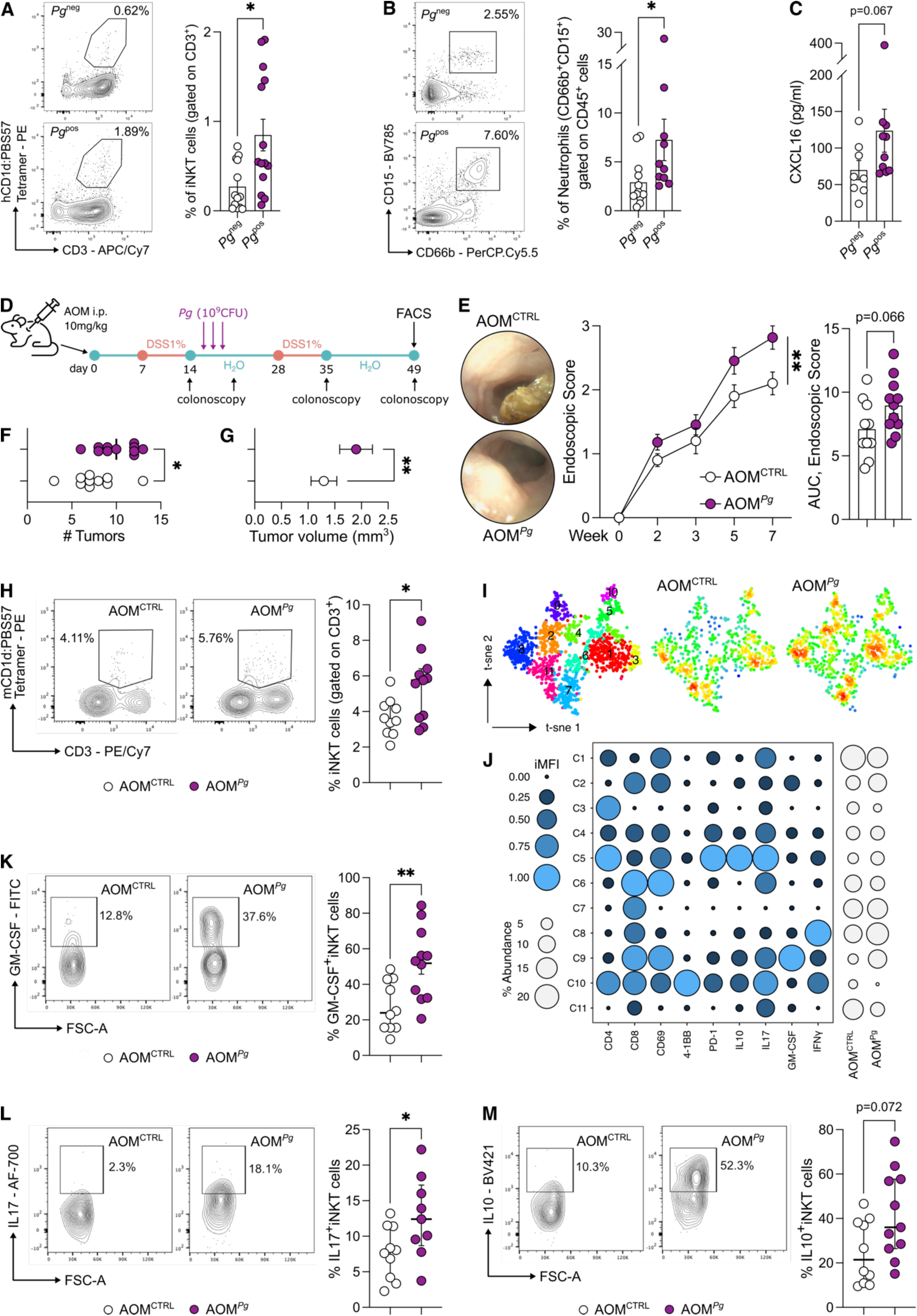
*P. gingivalis* promotes colorectal tumorigenesis by modulating iNKT cell functions. **A)** Frequency of tumour-infiltrating iNKT cells and **B)** tumour-associated neutrophils in CRC patients positive (*Pg*^pos^) or negative (*Pg*^neg^) for *P. gingivalis* in their mucosa-associated microbiota with representative dot plots. **C)** CXCL16 concentration from tissue lysates. **(D**) Schematic representation of the AOM-DSS experimental plan. **E)** Tumour endoscopic score, AUC and representative endoscopic pictures, **F)** number and **G)** volume of tumours from AOM-DSS treated C57BL/6 animals orally gavaged with PBS (AOM^CTRL^) or 10^9^ CFUs of *P. gingivalis* (AOM*^Pg^*). **H)** Frequency of tumour-infiltrating iNKT cells in AOM^CTRL^ and AOM*^Pg^* C57BL/6 mice with representative plots. **I)** t-SNE map of iNKT cells based on Phenograph metaclustering analysis of AOM^CTRL^ and AOM*^Pg^* tumour samples. **J)** Balloon plot of the scaled integrated Mean Fluorescent Intensity (iMFI) of Phenograph clusters generated in panel J. **K-M)** Frequency of tumour-infiltrating **K)** GM-CSF^+^ **L)** IL17^+^ and **M)** IL10^+^ iNKT cells in AOM^CTRL^ and AOM*^Pg^* C57BL/6 mice with representative plots. Data points (n=10, AOM^CTRL^; n=11, AOM*^Pg^*) from two pooled independent experiments representative of at least three. P < 0.05 (*), P < 0.01 (**); Mann-Whitney tests.

The upregulation of *IL17A*, *CD274* (*PD-L1*), and *CCL20* in *Pg*^positive^ patients suggested that iNKT cells might accumulate within an immunosuppressive TME, where the cytotoxic functions akin to Th1-like responses are dampened. This hypothesis is further suggested by the diminished expression of *TBX21*, encoding for the Th1 transcription factor T-bet (Supplementary Figure S1A). Moreover, we found in *Pg*^positive^ patients a significantly higher concentration of intratumour CXCL16, a key chemokine responsible for recruiting iNKT cells (*16*) (Figure 1C).

Next, we evaluated the contribution of *Pg* to CRC pathogenesis by using the chemical azoxymethane-dextran sodium sulphate (AOM-DSS) model of colitis-associated CRC (Figure 1D). *Pg*-treated C57BL/6 (B6) mice (AOM*^Pg^*) showed an increased tumour burden compared to control tumour-bearing mice (Figure 1E-G). A multidimensional immunophenotyping of T cells by Phenograph unsupervised clustering showed a different distribution of CD3^+^T cell density between AOM*^Pg^* vs AOM^CTRL^ B6 mice (Supplementary Figure 1B-C) although the frequencies of tumour-infiltrating CD4^+^ and CD8^+^ T cells as well as of NK cells did not differ by manual gating FACS analysis (Supplementary Figure S1D). In contrast, iNKT cells were significantly enriched in AOM*^Pg^* samples (Figure 1H). The metaclustering analysis of the iNKT cell cluster C2 (Supplementary Figure S1C) showed that tumour-infiltrating iNKT cells in AOM*^Pg^* mice are characterized by an overall increased expression of IL17, GM-CSF and IL10 in CD8^+^iNKT cells (C9; p=0.08, Mann-Whitney U test) (Figure 1I-J). We further confirmed the overall higher expression of GM-CSF, IL17 and IL10 in iNKT cells by manual gating FACS analysis (Figure 1K-M).

To demonstrate that *Pg* promotes CRC by modulating iNKT cell functions, we induced tumorigenesis in iNKT cell deficient *Traj18*^-/-^ mice (*Jα18*^-/-^). The results obtained underscore the essential role of iNKT cells in *Pg*-mediated CRC tumorigenesis. Specifically, tumour-bearing *Jα18*^-/-^ mice treated with *Pg* exhibited a comparable tumour burden to control AOM-DSS *Jα18*^-/-^ animals (Figure 2A-C). Intriguingly, frequencies and phenotypes of NK cells and conventional T cells remained largely unchanged in *Jα18*^-/-^ mice irrespective of *Pg* treatment (Supplementary Figure S1E), further supporting the notion of a specific interplay between *Pg* and iNKT cells.

**Figure 2:**
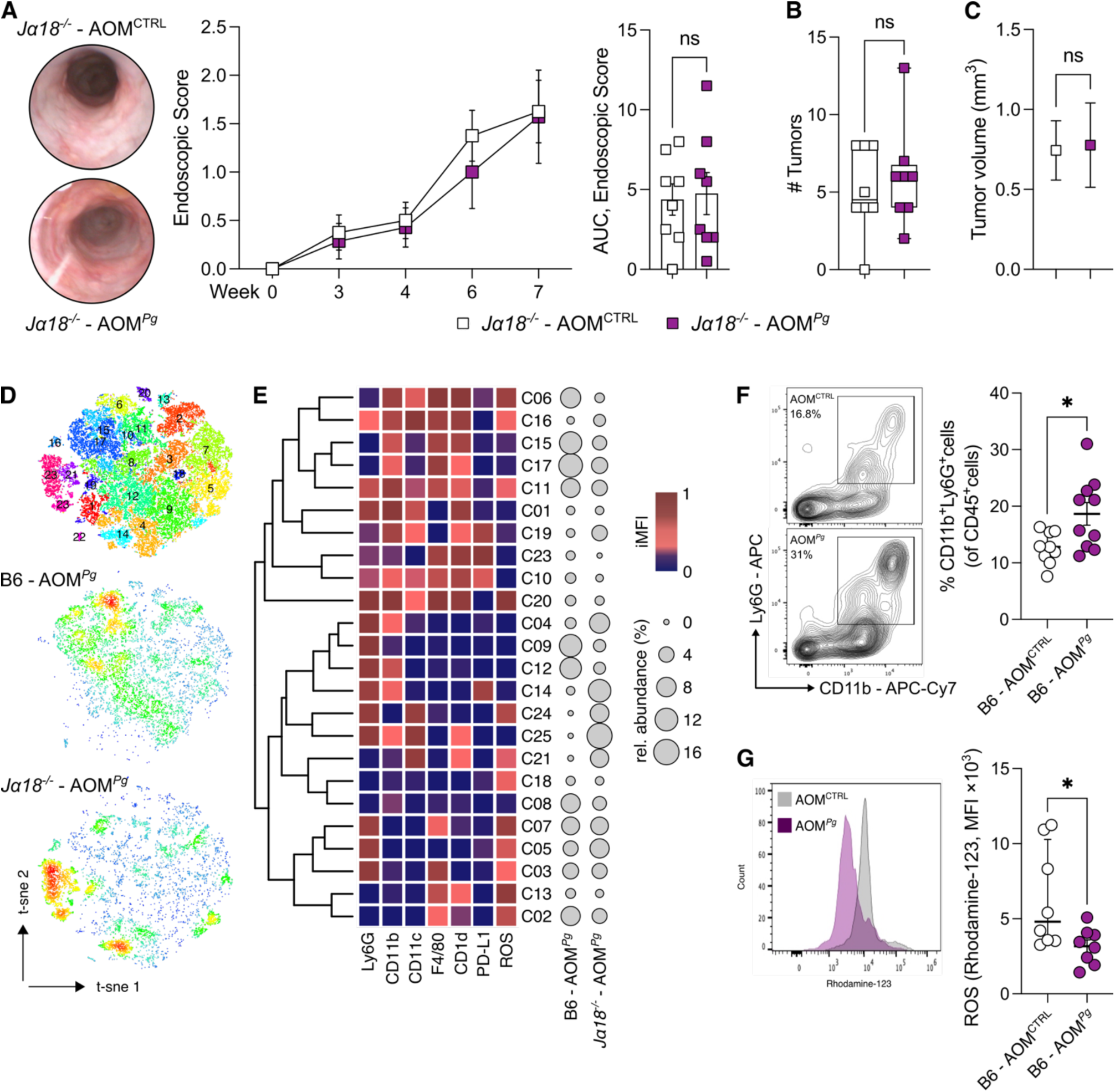
iNKT cells are essential to promote *P. gingivalis*-elicited colorectal tumorigenesis. **A)** Tumour endoscopic score, AUC and representative endoscopic pictures, **B)** number and **C)** volume of tumours from AOM-DSS treated *Traj18*^-/-^ mice (*Jα18*^-/-^) orally gavaged with PBS (AOM^CTRL^) or 10^9^ CFUs of *P. gingivalis* (AOM*^Pg^*). **D)** t-SNE map of intratumour myeloid cells based on Phenograph clustering from AOM-DSS, *P. gingivalis* treated C57BL/6 (B6 - AOM*^Pg^*) and *Traj18*^-/-^ mice (*Jα18*^-/-^ - AOM*^Pg^*). **E)** Heatmap of scaled integrated MFI data from Phenograph clustering analysis; relative abundance of the identified clusters is also shown. **F)** Frequency of CD11b^+^Ly6G^+^ and **G)** respiratory burst quantification of tumour-associated neutrophils from AOM-DSS treated C57BL/6 mice orally gavaged with PBS (AOM^CTRL^) or 10^9^ CFUs of *P. gingivalis* (AOM*^Pg^*), with representative dot plots. Data points (n=8-10) from two pooled independent experiments representative of at least three. P < 0.05 (*), P < 0.01 (**), P < 0.001(***); Mann-Whitney tests.

Then, we performed a multidimensional Phenograph analysis of tumour-infiltrating myeloid cells in tumour-bearing B6 and *Jα18*^-/-^ mice treated with *Pg* (Figure 2D-E) to investigate how *Pg* may condition the composition of myeloid cell populations through iNKT cells. *Pg* specifically induced the iNKT cell-dependent enrichment of CD11b^+^monocytic cells (C08; p=0.03, One-way ANOVA), F4/80^+^CD1d^+^ activated (ROS^+^) macrophages (C02; p=0.06, One-way ANOVA) and Ly6G^+^CD11b^+^neutrophils (C12; p=0.08, One-way ANOVA) (Figure 2E). In agreement with our previous study (*12*), we observed the enrichment of TANs in *Pg*-treated B6 mice compared to untreated AOM-DSS animals (Figure 2F). This enrichment is accompanied by TANs reduced respiratory burst capacity (Figure 2G), suggesting a diminished cytotoxic potential (*17*).

In summary, these data show that iNKT cells manifest a protumour functional phenotype modulating the mononuclear-phagocyte cell landscape through interaction with *Pg*.

### *Porphyromonas gingivalis* impairs iNKT cell cytotoxicity while promoting iNKT cell-mediated recruitment of TANs

To unravel the mechanisms governing the interaction between *Pg* and iNKT cells, we performed a series of experiments involving the priming of intestinal and circulating human iNKT cell lines (*18, 19*) with monocyte-derived dendritic cells (moDC) loaded with *Pg*. Subsequently, we conducted a range of *in vitro* functional assays and RNA sequencing on iNKT cells (Figure 3A). iNKT cells primed with *Pg* display a pronounced enrichment in genes associated with neutrophil chemotaxis, marked by the upregulation of key chemokines such *as CXCL1, CXCL2, CXCL5, CXCL8, CCL2*, and *CCL4* (Figure 3B-C). This finding strongly suggests that *Pg* can foster the iNKT cell-mediated recruitment of neutrophils. To validate this, we performed a migration assay that confirmed the capacity of *Pg*-primed iNKT cells to induce neutrophil chemotaxis (Figure 3D). Furthermore, iNKT cell activation induced by *Pg* led to a reduction in the respiratory burst capability of neutrophils (Figure 3E) coupled with an elevated expression of the immunosuppressive marker PD-L1 (Figure 3F) and an increase in neutrophil survival rates (Figure 3G).

**Figure 3:**
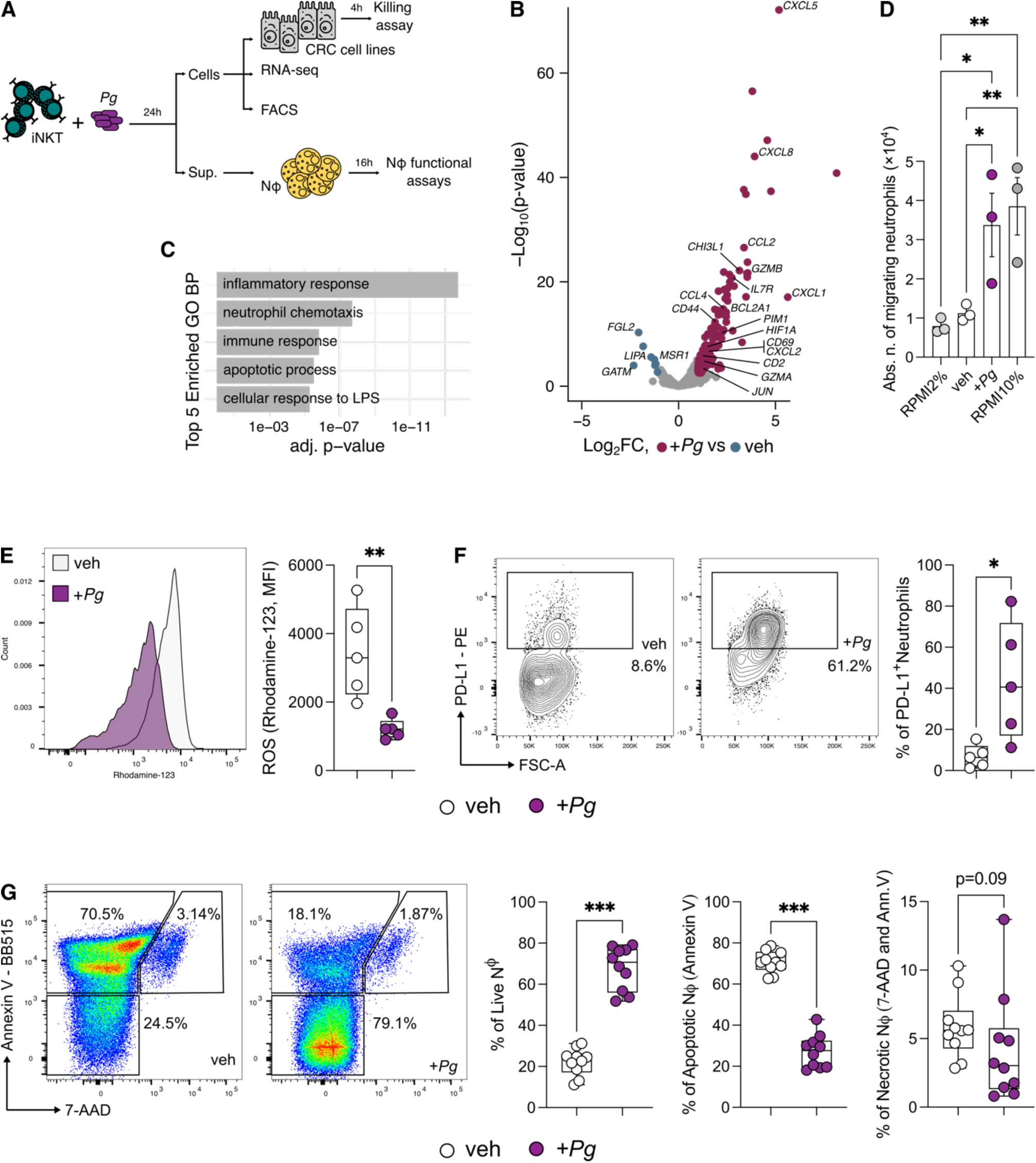
*P. gingivalis* promotes iNKT cell-mediated recruitment of TANs. **A)** Schematic representation of the experimental plan. **B)** Volcano plot representing the differentially expressed genes (DEGs) of *Pg-*primed iNKT cells (+*Pg*) vs basal activated iNKT cells (veh); the volcano plot shows for each gene (dots) the differential expression (log_2_fold-change [log_2_FC]) and its associated statistical significance (log_10_p-value). Dots indicate those genes with an FDR-corrected p < 0.05 and log_2_FC > |1|. **C)** Gene Ontology (GO) analysis of differentially expressed genes (Bonferroni-corrected p < 0.05 and log_2_FC > 2). **D)** Absolute numbers of migrating neutrophils upon exposure to RPMI+2%FBS (negative control), RPMI+10%FBS (positive control), basal activated (veh) and *Pg*-primed (+*Pg*) iNKT cell supernatants. Results are representative of three (n=3) independent experiments. P < 0.05 (*), P < 0.01 (**), One-Way ANOVA. **E)** Respiratory burst assay quantification and **F)** frequency of PD-L1^+^ cells from neutrophils exposed to the culture supernatants of *Pg-*primed (+*Pg*) and basal activated (veh) iNKT cells with representative plots. **G)** Representative plots and frequency of live (7-AAD^-^, Annexin V^-^), necrotic (7-AAD^+^ and Annexin V^+^) and apoptotic (7-AAD^-^, Annexin V^+^) neutrophils exposed to the culture supernatants of *Pg-*primed (+*Pg*) and basal activated (veh) iNKT cells. Data are representative of at least three independent experiments. P < 0.05 (*), P < 0.01 (**), P < 0.001(***), Mann-Whitney tests.

From a phenotypic perspective, *Pg* induced a Th17-like profile in iNKT cells, characterized by the expression of IL17, GM-CSF, and IL10 (Figure 4A-C). Functionally, *Pg* abrogated the cytotoxic capabilities of iNKT cells against colon adenocarcinoma cell lines (Figure 4D) because of the reduced release of perforin (PFN) and granzyme B (GMZB) (Figure 4E-F) and impaired lytic degranulation as evidenced by decreased CD107a expression (Figure 4G).

**Figure 4:**
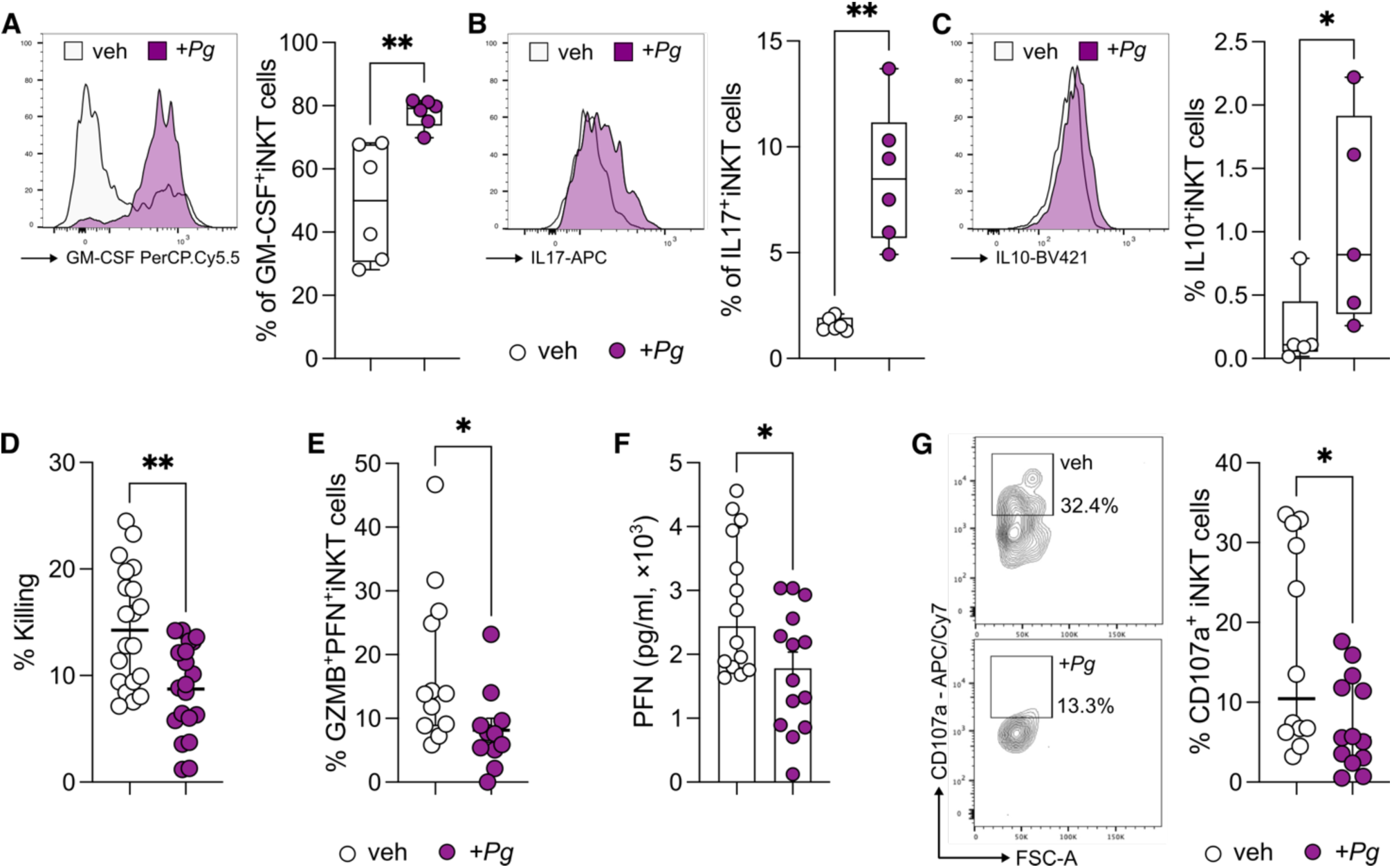
*P. gingivalis* impairs iNKT cell cytotoxicity. **A-C)** Frequency of **A)** GM-CSF^+^ **B)** IL17^+^ and **C)** IL10^+^ iNKT cells upon stimulation with *P. gingivalis*. **D)** Percentage of killed tumour cells by *Pg-*primed (+*Pg*) and basal activated (veh) iNKT cells. **E)** Frequency of GZMB^+^PFN^+^iNKT cells. **F)** Perforin concentration in the culture supernatant of *Pg-*primed (+*Pg*) and basal activated (veh) iNKT cells. **G)** Frequency of CD107a^+^iNKT cells with representative plots. Data are representative of at least three independent experiments. P < 0.05 (*), P < 0.01 (**), P < 0.001(***), Mann-Whitney tests.

Collectively, these findings suggest that *Pg* has the potential to foster CRC tumorigenesis by compromising the cytotoxic functions of iNKT cells.

### *Porphyromonas gingivalis* impairs iNKT cell cytotoxicity through CHI3L1

The RNA-seq analysis of *Pg*-primed iNKT cells revealed the upregulation of *CHI3L1* (Figure 3B). CHI3L1 is known to be a proinflammatory protein (*20*) that exerts its influence on the cytotoxic machinery of NK cells, resulting in the inhibition of their killing functions (*21*). Thus, we hypothesized that *Pg* might undermine iNKT cell cytotoxicity through a similar mechanism. In agreement with its mRNA expression, we measured a higher concentration of CHI3L1 in the culture supernatant of *Pg*-primed iNKT cells (Figure 5A).

**Figure 5:**
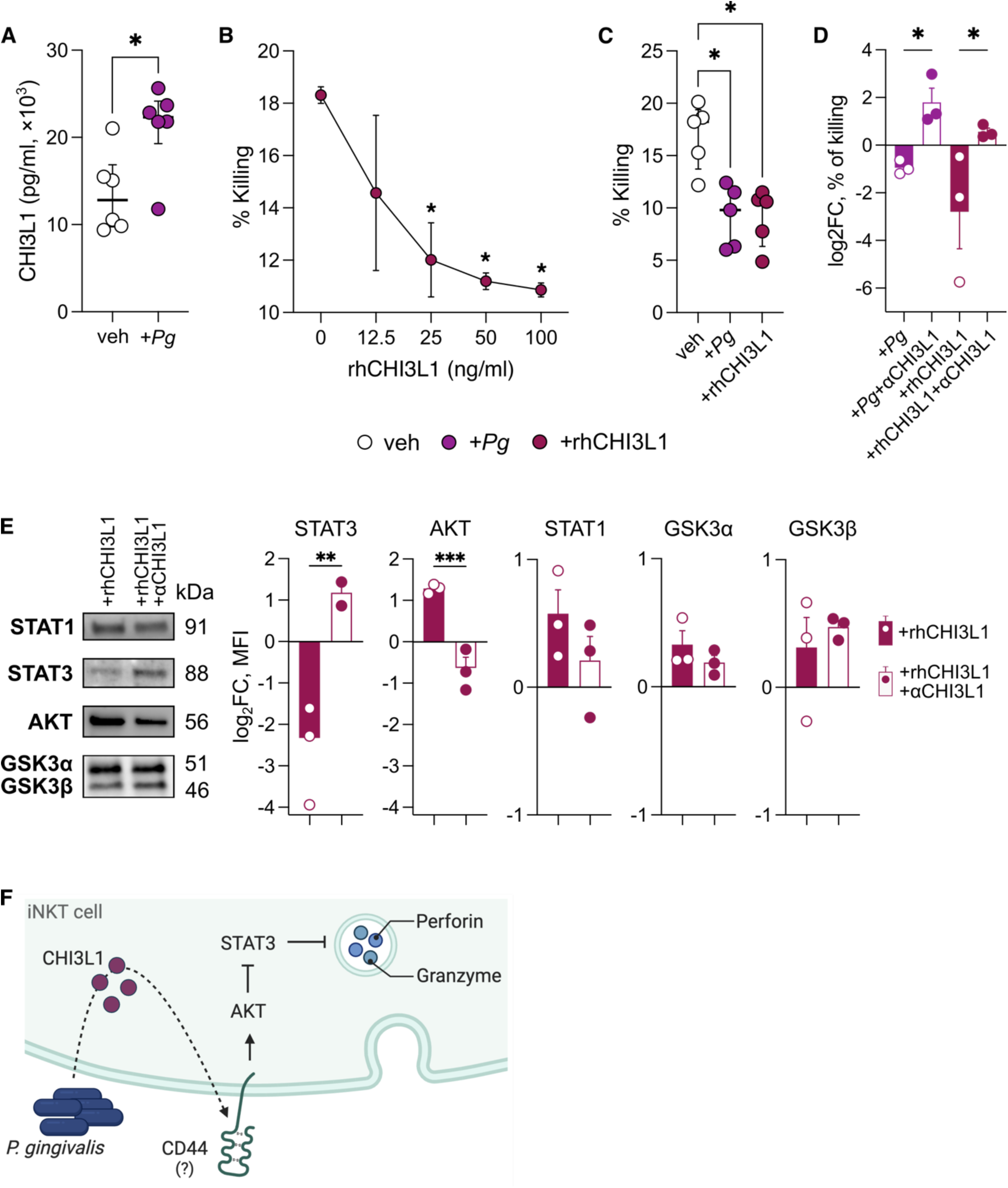
*P. gingivalis* exerts its effect on iNKT cells through CHI3L1. **A)** CHI3L1 concentration in the culture supernatant of *Pg-*primed (+*Pg*) and basal activated (veh) iNKT cells. **B)** Percentage of killed tumour cells by iNKT cells treated with increasing doses of rhCHI3L1. **C)** Percentage of killed tumour cells by iNKT cells basal activated (veh), primed with 10^9^ CFUs of *P. gingivalis* (+*Pg*) or treated with rhCHI3L1 (100ng/ml). **D)** Log_2_ fold-change of the percentage of killed tumour cells by iNKT cells primed with 10^9^ CFUs of *P. gingivalis* (+*Pg*) or treated with rhCHI3L1 (100ng/ml) upon neutralization with anti-CHI3L1 antibody (full bars); data have been normalized vs basal activated iNKT cells. **E)** Western blot analysis; MFI is expressed as log_2_ fold-change normalized vs basal activated iNKT cells. Data are representative of at least three independent experiments. P < 0.05 (*), P < 0.01 (**), P < 0.001(***); Mann-Whitney test and One-Way ANOVA. **F)** Proposed model; *Porphyromonas gingivalis* induces the expression and secretion of CHI3L1 in iNKT cells through the expression of an unidentified virulence factor (dotted arrow). The putative interaction with CD44, a known ligand of CHI3L1, leads to the activation of the PI3K/AKT pathway and repression of STAT3 signalling with the subsequent inhibition of cytotoxic degranulation.

In order to assess whether CHI3L1 directly affects iNKT cell cytotoxicity, we pre-treated human iNKT cell lines with varying concentrations of CHI3L1 before their co-incubation with target cells. Our findings indicate that CHI3L1 indeed impairs iNKT cell cytotoxicity in a dose-dependent manner (Figure 5B), mirroring the effects observed with *Pg* treatment (Figure 5C). To further investigate this aspect, we neutralized CHI3L1 using an anti-human CHI3L1 antibody on iNKT cells treated with both *Pg* and CHI3L1. This intervention successfully restored iNKT cell functions (Figure 5D) and STAT3 signalling (Figure 5E), a transcription factor critical to promote tumour-specific cytotoxic T cell development and effector functions in cancer (*22*). Collectively, these results highlight the potential of targeting CHI3L1 as a therapeutic strategy to restore iNKT cell activity.

## Discussion

Numerous studies have unveiled the capacity of the gut microbiome to regulate iNKT cell functions in health and disease (*12, 18, 23–25*). Nonetheless, the mechanisms by which gut microbes exert their influence on iNKT cells are not fully elucidated, and warrant further investigations for their safe use for adoptive cell therapies (*14*). The identification of such mechanisms assumes paramount importance given our recent findings in which we demonstrated that tumour-infiltrating iNKT cells bear unfavourable prognostic implications in human CRC because of the functional interaction with the periodontal pathogen *F. nucleatum* (*12*). Although iNKT cells are recognized as crucial components of anti-tumour immunity and their infiltration within tumour lesions is regarded as a positive prognostic factor (*26, 27*), in the presence of *F. nucleatum* these cells acquire a protumour Th17-like phenotype that facilitates the recruitment of immune suppressive TANs fostering CRC progression (*12*).

*P. gingivalis* is a keystone periodontal pathogen that is enriched in CRC and associated with poor overall and relapse free survival (*10, 11*). Here, we show that *P. gingivalis* increases the intratumour abundance of pro-inflammatory, yet immune suppressive iNKT cells in CRC patients and in *in vivo* models of CRC. Prior studies have indeed emphasized the elicitation of IL17- and IL10-mediated responses by *P. gingivalis* (*28–32*). *P. gingivalis* has the potential to enhance Th17 responses by upregulating the expression of key cytokines such as IL-6, IL-23, and IL-1β on myeloid cells, while simultaneously exerting a suppressive effect on IL-12 production (*31*). Remarkably, the involvement of gingipains and FimA, two significant *P. gingivalis* virulence factors (*32*), emerge as pivotal determinants of IL17 and IL10 production and the promotion of pathogenic responses in chronic inflammatory conditions (*28–30*). Moreover, *P. gingivalis* fosters an inflammatory response in CRC achieved through the recruitment of tumour-infiltrating CD11b^+^ myeloid cells and activation of the NLPR3 inflammasome (*11*). However, evidence regarding the mechanism by which *P. gingivalis* mediates the active recruitment of these myeloid cells within the TME is still lacking. Our study suggests that the *Pg*-mediated recruitment of myeloid cells in tumour lesions might be predominantly orchestrated by iNKT cells. Indeed, we demonstrated that iNKT cells elicit remarkable changes in the intratumour mononuclear-phagocyte cell landscape in response to *P. gingivalis* while the composition of conventional CD4^+^ and CD8^+^ T cells within the TME remains unaltered, as shown also in previous studies (*11*). *P. gingivalis* has also a distinguishing pathogenic trait in the impairment of iNKT cell cytotoxic functions, when compared to its oral counterpart *F. nucleatum.* Indeed, *P. gingivalis* appears to specifically interfere with the iNKT cell lytic granule machinery, since we observed a decreased secretion of perforin and reduced expression of the degranulation marker CD107a. The immune-mediated elimination of cancer cells depends on the lytic granule machinery of cytotoxic lymphocytes, including iNKT cells, CD8^+^ cytotoxic T lymphocytes and natural killer (NK) cells. Nevertheless, cancer cells can use a variety of evasion mechanisms to prevent these cells from killing them.

CHI3L1 protein is a mammalian member of the evolutionarily conserved chitinase protein family. Although its precise physiological function remains incompletely understood, aberrant expression of CHI3L1 is linked to the development of various human diseases, including cancer (*20*). CHI3L1 expression by T cells promotes lung metastasis by dampening antitumour Th1 responses (*33*). When acting on NK cells, a population that shares phenotypic and functional features with iNKT cells, CHI3L1 disrupts the proper alignment of the microtubule-organizing center and lytic granules at the immunological synapse (*21*). Our study reveals that *P. gingivalis* induces the production of CHI3L1 by iNKT cells, thereby diminishing their ability to eliminate CRC cells. Remarkably, neutralization of CHI3L1 is sufficient to restore iNKT cell activity, even in the presence of exogenous CHI3L1 supplementation. CHI3L1 can signal through CD44, one of its physiological receptors (*34–36*), which is upregulated in *Pg*-primed iNKT cells (Figure 3B). This signalling may act as an immune checkpoint to suppress iNKT cell activation via the PI3K/AKT pathway, a known repressor of STAT3 transcription (Figure 5F) (*22, 37–39*), but warrants further investigations. In conclusion, our data demonstrate that *P. gingivalis* fuels CRC progression by inducing iNKT cells to express CHI3L1, an immune checkpoint that can suppress iNKT cell cytotoxicity favouring host tumour immune evasion.

## Materials & Methods

### Human samples

Samples were collected with informed consent from patients (n = 32) diagnosed with colorectal adenocarcinoma between January 2017 and July 2022 undergoing surgical resection at IRCCS Policlinico Ospedale Maggiore, Milan, Italy, as approved by the Institutional Review Board (Milan, Area B) with permission number 566_2015.

### Tumour-associated microbiota isolation

The tumour-associated microbiota was obtained at the moment of surgery from tumour tissue by scraping. Handling, DNA extraction, sequencing and analysis was performed as previously described (*12*).

### Isolation of tumour-infiltrating cells

Tumour samples were taken transversally to collect both marginal and core tumour zone. Human lamina propria mononuclear cells (LPMCs) were isolated as previously described (*40*).

### Mice

C57BL/6 and B6(Cg)-Traj18tm1.1Kro/J (*Traj18*^-/-^) mice (*41*) (provided by G. Casorati and P. Dellabona, San Raffaele Scientific Institute) were housed and bred at the European Institute of Oncology (IEO) animal facility (Milan, Italy) in SPF conditions. Sample size was chosen based on previous experience. No sample exclusion criteria were applied. No method of randomization was used during group allocation, and investigators were not blinded. Age-matched male and female mice were used for experiments. Animal experimentation was approved by the Italian Ministry of Health (Auth. 10/21 and Auth. 1217/20) and by the animal welfare committee (OPBA) of the European Institute of Oncology (IEO), Italy.

### Porphyromonas gingivalis culture condition

*P. gingivalis* strain DSM 20709 (ATCC 33277) was maintained on Columbia agar supplemented with 5% sheep blood or in Columbia broth (Difco, Detroit, MI, USA) under anaerobic conditions at 37 °C. Columbia broth was supplemented with hemin at 5 μg·mL−1 and menadione at 1 μg·mL−1. Bacterial cell density was adjusted to 1 × 10^10^ CFU·mL−1 and heat-killed at 95°C for 15min before being stored at −80 °C until use in downstream experimentation.

### Animal experiments

7-8 weeks old mice were injected intraperitoneally with azoxymethane (AOM, Merck) dissolved in isotonic saline solution at a concentration of 10 mg/kg body weight. After 7 days, mice were given 1% (w/v) dextran sodium sulfate (DSS MW 40 kD; TdB Consultancy) in their drinking water for 7 days followed by 14 days of recovery. The cycles were repeated for a total of 2 DSS cycles, and mice sacrificed at day 49. After the first cycle of DSS treatment, during the recovery phase, mice were orally gavaged for 3 days with 200 μl of PBS (control) or 10^9^ CFUs (colony forming unit) suspension of *P. gingivalis* DSM 20709. The treatment schedule is shown in Figure 1D.

### Murine colonoscopy

Colonoscopy was performed weekly for tumour monitoring using the Coloview system (TP100 Karl Storz, Germany). Tumour endoscopic score has been quantified as previously described (*42*). During the endoscopic procedure mice were anesthetized with 3% isoflurane.

### Murine cells isolation

Single-cell suspensions were prepared from the colon of C57BL/6 and *Traj18*^-/-^ mice as previously described (*24*). Briefly, cells were isolated via incubation with 5 mM EDTA at 37°C for 30 min, followed by mechanical disruption with GentleMACS (Miltenyi Biotec). After filtration with 100-µm and 70-µm nylon strainers (BD), the LPMC were counted and stained for immunophenotyping.

### Cell lines

The different cell lines used in this work are listed in Supplementary Table S1. Human iNKT cell lines were generated as previously described (*18*). All cells were maintained in a humidified incubator with 95% air, 5% CO2 at 37°C.

### Pg-priming of iNKT cell

1 × 10^5^ monocyte derived dendritic cells (moDCs) were pulsed with heat-inactivated *P. gingivalis* (*Pg*) (4 × 10^5^ CFU) and co-cultured with iNKT cells (2 × 10^5^ cells) in RPMI-1640 supplemented with 10%FBS, Pen/Strep. After 24h, iNKT cell activation status was estimated by extracellular or intracellular staining.

### iNKT cell cytotoxicity assay

iNKT cell cytotoxicity toward the human CRC cell lines Colo205 and RKO (American Type Culture Collection, ATCC) was performed as previously described (*19*). In CHI3L1 neutralization experiments, anti-CHI3L1 (10 µg/ml, clone mYA, Millipore) were pre-incubated with rhCHI3L1 (R&D Systems) or *Pg*-primed iNKT cells for 24 hours at 37°C before performing the assay.

### Neutrophil isolation

Neutrophils were isolated from whole blood samples by dextran sedimentation (4% diluted in HBSS). Red blood cells were lysed using ACK lysis buffer (Life Technologies) and neutrophils separated with Percoll gradient.

### iNKT cell-Neutrophil co-culture assay

*P. gingivalis* primed-iNKT cells (2 × 10^5^ cells) were co-cultured with freshly isolated neutrophils in a 1:1 ratio, in RPMI-1640 supplemented with 10% FBS. After 24h cells were stained for the expression of extracellular marker and ROS production.

### Neutrophil migration assay

Freshly isolated neutrophils were seeded on top of a 3μm-pore transwell (SARSTEDT) in 200μl of RPMI-1640+2%FBS. 500μl of chemoattracting medium was added to the bottom of transwells and neutrophil migration was allowed for 4 hours at 37°C. RPMI-1640+10% FBS was used as positive control. The total number of cells on the bottom of the plate was stained and counted using the FACSCelesta flow cytometer (BD Biosciences, Franklin Lakes NJ, USA) with plate-acquisition mode and defined volumes.

### Neutrophil survival assay

Freshly isolated neutrophils were cultured with RPMI-1640+10% FBS supplemented with the culture supernatants (10%) from *Pg*-primed iNKT cells for 16h at 37°C. Cells were then stained with FITC Annexin V Apoptosis Detection Kit with 7-AAD (Biolegend) following manufacturer’s instruction and acquired at a FACS Celesta flow cytometer (BD Biosciences, Franklin Lakes NJ, USA).

### Respiratory Burst Assay

ROS production was quantified using the Neutrophil/Monocyte Respiratory Burst assay (Cayman) following manufacturer’s instructions.

### ELISA assay

Detection of Perforin and CHI3L1 in iNKT cell culture supernatants was performed using the Human Perforin Elisa Flex (Mabtech) and the Human CHI3L1 DuoSet ELISA (R&D systems) according to manufacturers’ instructions.

### Flow Cytometry

Cells were washed and stained with the combination of mAbs purchased from different vendors, as listed in Supplementary Table S1. iNKT cells were stained and identified using human or mouse CD1d:PBS57 Tetramer (NIH Tetramer core facility) diluted in PBS with 1% heat-inactivated FBS for 30 min at 4°C. For intracellular cytokine labelling cells were incubated for 3 h at 37°C in RPMI-1640+10% FBS with PMA (50ng/ml, Merck), Ionomycin (1μg/ml, Merck) and Brefeldin A (10 μg/ml, Merck). Before intracellular staining cells were fixed and permeabilized using Cytofix/Cytoperm (BD). Samples were analysed with a FACSCelesta flow cytometer (BD Biosciences, Franklin Lakes NJ, USA) and the FlowJo software (Version 10.8, TreeStar, Ashland, OR, USA). For the multidimensional analysis using t-SNE visualization and Phenograph clustering (*43*), FCS files were quality checked for live, singlets and antibody agglomerates and normalized to avoid batch effects. Multidimensional regression and clustering analysis were performed as previously described (*12*).

### Bulk RNA sequencing of human iNKT cells

Total RNA (from 1 × 10^6^ cells) was isolated with the RNeasy kit (Qiagen) and RNA quality was checked with the Agilent 2100 Bioanalyzer (Agilent Technologies). 0.5-1 μg were used to prepare libraries for RNA-seq with the Illumina TruSeq RNA Library Prep Kit v2 following the manufacturer’s instructions. RNA-seq libraries were then run on the Agilent 2100 Bioanalyzer (Agilent Technologies) for quantification and quality control and pair-end sequenced on the Illumina NovaSeq platform.

### RNA sequencing data analysis

RNA-seq reads were preprocessed using the FASTX-Toolkit tools. Quality control was performed using FastQC. Pipelines for primary analysis (filtering and alignment to the reference genome of the raw reads) and secondary analysis (expression quantification, differential gene expression) have been integrated and run in the HTS-flow system (*44*). Differentially expressed genes were identified using the Bioconductor Deseq2 package (*45*). P-values were False Discovery Rate corrected using the Benjamini-Hochberg procedure implemented in DESeq2. Functional enrichment analyses to determine Gene Ontology categories and KEGG pathways were performed using the DAVID Bioinformatics Resources (DAVID Knowledgebase v2022q2) (https://david.ncifcrf.gov) (*46*).

### Western Blot

Total protein extracts from iNKT cells were prepared as previously described (*47*), separated on Mini-PROTEAN TGX Stain-Free Precast Gels (4-15%, Bio-Rad Laboratories, Hercules, California, USA) to enhance transfer efficiency and detection of proteins with stain-free enabled imagers. Samples were then transferred on nitrocellulose membranes (Bio-Rad Laboratories, Hercules, California, USA) and incubated overnight at 4°C with the following primary antibodies: STAT-1 (1:600, E-Ab-32977, Elabscience, Houston, Texas, USA); STAT-3 (1:600, E-Ab-40131, Elabscience, Houston, Texas, USA); GSK-3ab (1:500, sc-7291, Santa Cruz, Dallas, Texas, USA); Akt 1/2/3 (1:500, Ab-179463, Abcam Cambridge, United Kingdom). Membranes were detected with peroxidase conjugated secondary antibodies (Agilent Technologies, California, USA) and developed by ECL (Amersham Biosciences, United Kingdom). Image Lab Software from Bio-Rad was used to analyse band intensity.

## Authors contribution

FF and FS conceived the study. FF, FS, ADB and GL designed the experiments. GL, ADB, FP, CA, AB and AF performed the experiments. FF and FS supervised the experiments. FC, MV, LB, YT, GP and MR contributed with reagents and resources. GL performed multidimensional FACS data analysis. FS performed RNAseq data analyses. FS wrote the first draft of the manuscript. All authors reviewed and critically edited the manuscript. Both GL and ADB contributed equally and have the right to list their name first in their CV. All authors contributed to the article and approved the submitted version.

## Acknowledgments

We thank the IEO Animal Facility for the excellent animal husbandry and the NIH Tetramer Facility for providing human and murine CD1d:PBS57 tetramers. We are grateful to the équipe of the General and Emergency Surgery Unit, Ospedale Maggiore Policlinico, Milano for their tireless work. We thank Prof. Paolo Dellabona and Prof. Giulia Casorati for providing the *Traj18*^-/-^ mice. We thank Maria Rita Giuffrè, Valentina Pasquale, Elisa Cirrincione, Luca Iachini, Luana Tripodi and Monica Molinaro for the assistance with the experiments. Panels 1D and 3A were created using icons from the Noun Project (https://thenounproject.com/). Panel 5F was created with BioRender (https://biorender.com/).

---

**Supplementary Figure 1:**
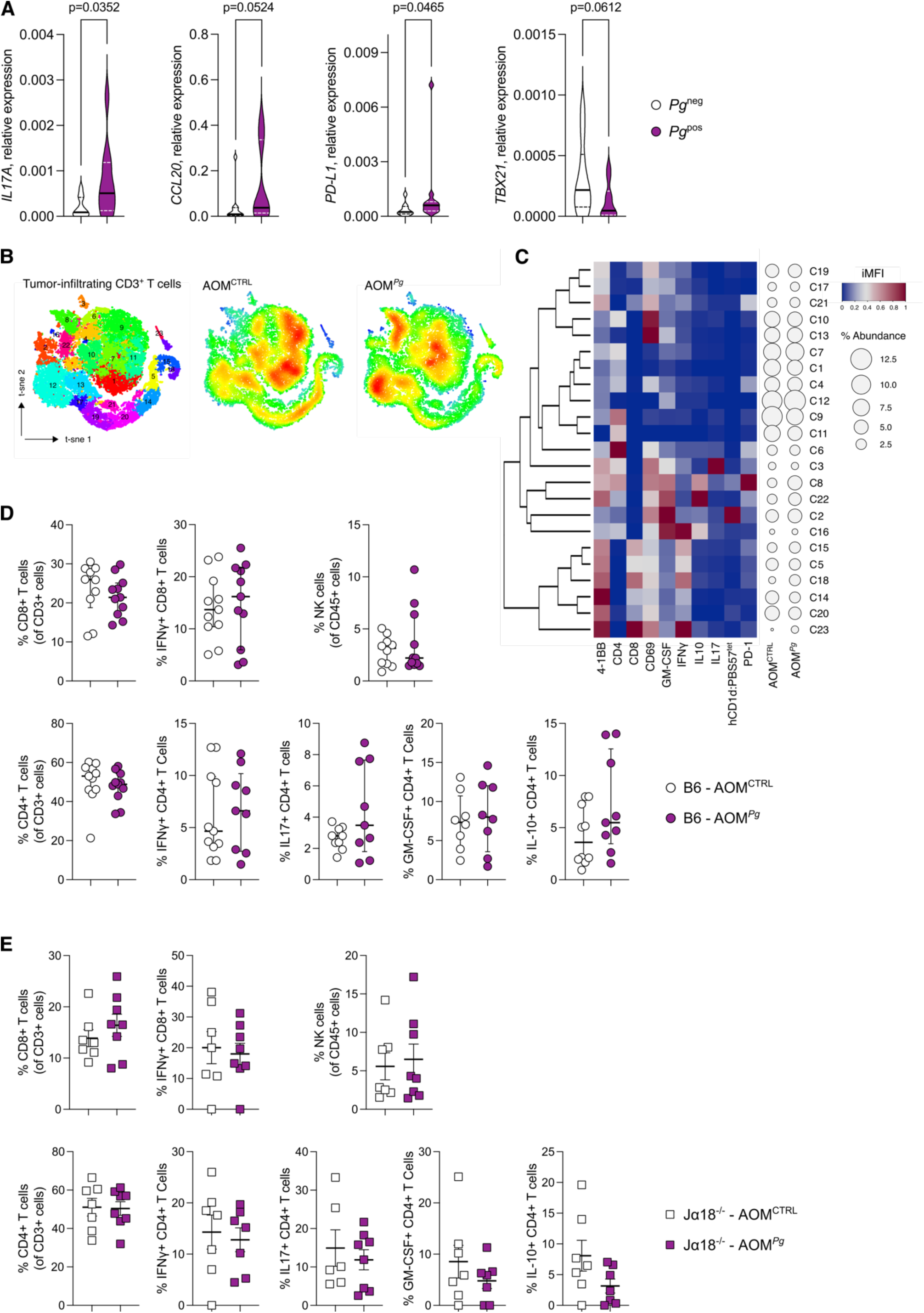
**A)** Tumour mRNA expression of *IL17A*, *CCL20*, *CD274 (PD-L1)* and *TBX21* in CRC patients harboring *P. gingivalis* in their mucosa-associated microbiota (*Pg*^pos^) **B)** t-SNE map of of CD3^+^T cells based on Phenograph clustering analysis of AOM^CTRL^ and AOM*^Pg^* tumour samples from B6 mice. **C)** Heatmap of scaled integrated MFI data from Phenograph clustering analysis; relative abundance of identified clusters is also shown. D-E) Frequency of NK cells, CD4^+^ and CD8^+^T cell in **D)** C57BL/6 and **E)** *Traj18*^-/-^ mice.

**Supplementary Table S1:** Materials and Resources.

| Reagent or Resource | Source | Identifier |
| --- | --- | --- |
| **Antibodies** |  |  |
| Anti-human CD274 (PDL1) PE conjugated | Biolegend | Cat.#:393608;Clone:MIH2;RRID:AB_2749925 |
| Anti-human CD274 (PDL1) PE-Cy7 conjugated | BD Biosciences | Cat.#:558017;Clone:MIH1;RRID:AB_396986 |
| Anti-human CD3 APC-Cy7 conjugated | TONBO | Cat.#:25-0038-T100;Clone:UCHT1 |
| Anti-human CD3 PE-Cy7 conjugated | TONBO | Cat.#:60-0038-T100;Clone:UCHT1 |
| Anti-human CD45 BV510 conjugated | BD Biosciences | Cat.#:563204;Clone:HI30;RRID:AB_2738067 |
| Anti-human CD66b PerCP-Cy5.5 conjugated | BD Biosciences | Cat.#:562254;Clone:G10F5;RRID:AB_11154419 |
| Anti-human GM-CSF PerCP-Cy5.5 conjugated | Biolegend | Cat.#:502312;Clone:BVD2-21C11;RRID:AB_11147946 |
| Anti-human IFN-γ FITC conjugated | BD Biosciences | Cat.#:554551;Clone:4S.B3;RRID:AB_395473 |
| Anti-human IL17A APC conjugated | ThermoFisher Scientific | Cat.#:11-7179-42;Clone:eBio64DEC17;RRID:AB_1582221 |
| Anti-human IL10 PerCP-eFluor710 conjugated | ThermoFisher Scientific | Cat.# 46-7108-42;Clone:JES3-9D7;RRID:AB_2573833 |
| Anti-Chitinase-3-like protein 1 | Millipore | Cat.#MABC196;Clone:mAY;RRID:AB_2891310) |
| Human CD1d:PBS57 Tet-PE conjugated | Gift from NIH Tet facility | N/A |
| Human CD1d:PE conjugated | Gift from NIH Tet facility | N/A |
| Anti-mouse CD11b APC-Cy7 conjugated | TONBO | Cat.#:25-0112-U100;Clone:M1/70 |
| Anti-mouse CD274 (PDL1) BV786 conjugated | BD Biosciences | Cat.#:741014;Clone:MIH5;RRID:AB_2740636 |
| Anti-mouse CD279 (PD1) APC conjugated | Biolegend | Cat.#:562671;Clone:J43;RRID:AB_2737712 |
| Anti-mouse CD3 FITC conjugated | Biolegend | Cat.#:100204;Clone:17A2;RRID:AB_312661 |
| Anti-mouse CD3 PE-Cy7 conjugated | Biolegend | Cat.#:100220;Clone:17A2;RRID:AB_1732057 |
| Anti-mouse CD4 BV650 conjugated | Biolegend | Cat.#:100545;Clone:RM4-5;RRID:AB_11126142 |
| Anti-mouse CD4 PE/Dazzle594 conjugated | Biolegend | Cat.#:100455;Clone:GK1.5;RRID:AB_2565844 |
| Anti-mouse CD45 Alexa Fluor 700 conjugated | Biolegend | Cat.#:103127;Clone:30-F11;RRID:AB_493714 |
| Anti-mouse CD45.2 APC-Cy7 conjugated | Biolegend | Cat.#:109824;Clone:104;RRID:AB_830789 |
| Anti-mouse CD45.2 BV510 conjugated | Biolegend | Cat.#:109838;Clone:104;RRID:AB_2650900 |
| Anti-mouse CD8a APC-Cy7 conjugated | BD Biosciences | Cat.#:557654;Clone:53-6.7;RRID:AB_396769 |
| Anti-mouse CD8a Super Bright 600 conjugated | ThermoFisher Scientific | Cat.#:63-0081-82;Clone:53-6.7;RRID:AB_2637163 |
| Anti-mouse F4/80 PE conjugated | TONBO | Cat.#:50-4801-U025;Clone:BM8.1 |
| Anti-mouse GM-CSF FITC conjugated | Biolegend | Cat.#:505404;Clone:MIP1-22E9;RRID:AB_315380 |
| Anti-mouse IL10 PE conjugated | Biolegend | Cat.#:554467;Clone:JES5-16E3;RRID:AB_395412 |
| Anti-mouse IL17A Alexa Fluor-700 conjugated | Biolegend | Cat.#:506914;Clone:TC11-18H10.1;RRID:AB_536016 |
| Anti-mouse IL17A BV605 conjugated | BD Biosciences | Cat.#:564169;Clone:TC11-18H10 |
| Anti-mouse Ly6G APC conjugated | TONBO | Cat.#:20-1276-U025;Clone:1A8 |
| Mouse CD1d:BV421 conjugated | Gift from NIH Tet facility | N/A |
| Mouse CD1d:PE conjugated | Gift from NIH Tet facility | N/A |
| Mouse CD1d:PBS57 Tet-BV421 conjugated | Gift from NIH Tet facility | N/A |
| Mouse CD1d:PBS57 Tet-PE conjugated | Gift from NIH Tet facility | N/A |
| **Chemicals, Peptides and Recombinant proteins** |  |  |
| ACK Lysis buffer | Life Technologies | Cat.#:A1049201 |
| Agar powder | Sigma-Aldrich | Cat.#:05040-250G |
| Azoxymethane | Merck Life Science | Cat.#:A5486 |
| Brefeldin A - 5 Mg | Merck Life Science | Cat.#:B7651-5MG |
| Collagenase D 2,5G from Clostridium histolyticum | Sigma-Aldrich | Cat.#:11088882001 |
| Columbia Agar + 5% Sheep Blood | ThermoFisher Scientific | Cat.#:PB5039A |
| Dextran from *Leuconostoc spp.* | Merck Life Science | Cat.#:31392-50G |
| Dextran Sulphate Sodium | Tdb Consultancy Ab | Cat.#:DB001-500G |
| DI-Dithiothreitol | Microtech S.R.L. | Cat.#:D0632-25G |
| DNA ladder | New England Biolabs | Cat.#:N3231L/2L |
| Dulbecco's Phosphate Buffered Saline | Microtech S.R.L. | Cat.#:TL1006-500ML |
| Easyscript Plus cDNA synthesis Kit | abm | Cat.#:G236 |
| Eurogold trifastII nucleic acid isolation reagent | Euroclone | Cat.#:EMR517200 |
| Fetal Bovine Serum | Microtech S.R.L. | Cat.#:RM10432 |
| Ficoll Paque Plus | Merck Life Science | Cat.#:GE17-1440-03 |
| Fixation/Permeabilization Solution Kit | BD Biosciences | Cat.#:554714 |
| Gentamicin | Himedia | Cat.#:A005-100ML |
| GentleMACS C Tubes | Miltenyi Biotec | Cat.#:130-096-334 |
| Hanks' Balanced Salt solution | Merck Life Science | Cat.#:H6648 |
| Hanks' Balanced Salt solution with Phenol Red | Himedia | Cat.#:TL1003-500ML |
| Human serum | Merck Life Science | Cat.#:H4522-100ML |
| Ionomycin calcium salt from *Streptomyces conglobatus* | Merck Life Science | Cat.#:I0634 |
| LS column | Miltenyi Biotec | Cat.#:130-042-401 |
| Maxisorp nunc immuno plate | ThermoFisher Scientific | Cat.#:M9410-1CS |
| MEM NEAA (100x) | ThermoFisher Scientific | Cat.#:11140-035 |
| Minimum Essential Medium Eagle | ThermoFisher Scientific | Cat.#:41090036 |
| Oxoid AnaeroGen 2.5L Sachet | ThermoFisher Scientific | Cat.#:AN0025A |
| Penicillin/Streptomycin | Microtech S.R.L. | Cat.#:A001-100ML |
| Percoll Plus | Merck Life Science | Cat.#:17-5445-01 |
| PMA (Phorbol 12-myristate 13-acetate) | Merck Life Science | Cat.#:P1585 |
| Proleukin, human IL2 recombinant protein | Novartis | Cat.#:27131010 |
| Qubit RNA High Sensitivity Assay Kit | ThermoFisher Scientific | Cat.#:Q32852 |
| Recombinant Human Chitinase 3-like 1 | R&D systems | Cat.#:2599-CH-050 |
| Remel PHA Purified | ThermoFisher Scientific | Cat.#:R30852801 |
| RNAprotect Cell Reagent | QIAGEN | Cat.#:76526 |
| RNaseZAP | Merck Life Science | Cat.#:R2020 |
| RPMI medium 1640 (1x) | Microtech S.R.L | Cat. #:AL028A-500 |
| Sodium Bicarbonate | Sigma-Aldrich | Cat. #:S6297-250G |
| Sodium Carbonate | Sigma-Aldrich | Cat. #:S7795-500G |
| Sodium Pyruvate 100mM | ThermoFisher Scientific | Cat. #:11360-039 |
| Streptavidin-HRP | Mabtech | Cat. #:3310-9 |
| Sulfuric Acid | Sigma-Aldrich | Cat. #:25810-5 |
| SYBR™ Safe DNA Gel Stain | ThermoFisher Scientific | Cat. #:S33102 |
| TC-insert 24 well PET 3um | SARSTEDT | Cat. #:83.3932.300 |
| TMB Chromogen Solution | ThermoFisher Scientific | Cat. #:2023 |
| Tris-EDTA buffer solution | Merck Life Science | Cat. #:T9285 |
| Trypsin-EDTA | ThermoFisher Scientific | Cat. #:25300-054 |
| **Bacterial strains** |  |  |
| *Porphyromonas gingivalis* | DSMZ | DSM20709 (ATCC33277) |
| **Critical commercial assay** |  |  |
| Cytotoxicity Ldh Assay Kit | Microtech S.R.L. | Cat. #:CK12-20 |
| Direct-zol Rna Miniprep Plus Kit | Zymo research | Cat. #:ZYR2072 |
| FITC Annexin V Apoptosis Detection Kit with 7-AAD | Biolegend | Cat. #:640922 |
| Human CHI3L1 DuoSet ELISA | R&D systems | Cat. #:DY2599 |
| Human Perforin ELISA Flex | Mabtech | Cat. #3465-1H-6(20) |
| Neutrophil/Monocyte Respiratory Burst Assay Kit | Cayman chemical | Cat. #:601130 |
| **Laboratory animals** |  |  |
| B6(Cg)-Traj18tm1.1Kro/J; TRAJ18-/- | Gift from P. Della Bona | JAX stock:030524;RRID:IMSR_JAX:030524 |
| C57BL/6 | The Jackson Lab | JAX stock:000664;RRID:IMSR_JAX:000664 |
| **Cell lines** |  |  |
| RKO | ATCC | CRL-2577™ |
| COLO205 | ATCC | CCL-222™ |
| PB5 | In house; (peripheral blood, healthy donor) | (Burrello et al., 2019; Díaz-Basabe et al., 2021) |
| PB6 | In house; (peripheral blood, healthy donor) | (Burrello et al., 2019; Díaz-Basabe et al., 2021) |
| NUN | In house; (colon, UC patient) | (Burrello et al., 2019; Díaz-Basabe et al., 2021) |
| CD2 | In house; (colon, CD patient) | (Burrello et al., 2019; Díaz-Basabe et al., 2021) |
| CD3 | In house; (colon, CD patient) | (Burrello et al., 2019; Díaz-Basabe et al., 2021) |
| **Software and Algorithms** |  |  |
| FACSDiva | BD bioscience | https://www.bdbiosciences.com |
| FlowJo v10.8 | BD bioscience | https://www.flojo.com/ |
| GraphPad Prism v9.5.2 | Graphpad Software | https://www.graphpad.com |
| R package (Biobase) | (Huber W et al., 2015) | https://github.com/Bioconductor/Biobase |
| R package (cytofkit) | (Chen H et al., 2016) | https://github.com/JinmiaoChenLab/cytofkit |
| R package (flowCore) | (Ellis B et al., 2022) | https://github.com/RGLab/flowCore |
| R package (flowStats) | (Hahne F et al., 2022) | https://github.com/RGLab/flowStats |
| R package (flowViz) | (Ellis B et al., 2022) | https://github.com/RGLab/flowViz |
| R package (Tidyverse and ggplot2) | (Wickham et al., 2019) | https://cloud.r-project.org/package=ggplot2 |
| R studio v4.1.2 & v3.6.2 | The R Foundation | https://cran.r-project.org/bin/macosx/ |
| DAVID Knowledgebase v2022q2 | (Sherman BT et al., 2022) | https://david.ncifcrf.gov |
| HTS-flow (High-Throughput Sequencing flow) | (Bianchi V et al., 2016) | https://github.com/arnaudceol/htsflow |
| R package "DESeq2" | (Love MI et al., 2014) | https://bioconductor.org/packages/release/bioc/html/DESeq2.html |

## References

1. S. V. S. Deo, J. Sharma, S. Kumar, GLOBOCAN 2020 Report on Global Cancer Burden: Challenges and Opportunities for Surgical Oncologists. Ann Surg Oncol 29, 6497–6500 (2022).

2. M. Schmitt, F. R. Greten, The inflammatory pathogenesis of colorectal cancer. Nat Rev Immunol 21, 653–667 (2021).

3. J. Chen, E. Pitmon, K. Wang, Microbiome, inflammation and colorectal cancer. Semin Immunol 32, 43–53 (2017).

4. D. Ternes et al., Microbiome in Colorectal Cancer: How to Get from Meta-omics to Mechanism? Trends Microbiol 28, 401–423 (2020).

5. G. Hajishengallis, Periodontitis: from microbial immune subversion to systemic inflammation. Nat Rev Immunol 15, 30–44 (2015).

6. N. Bostanci, G. N. Belibasakis, Porphyromonas gingivalis: an invasive and evasive opportunistic oral pathogen. FEMS Microbiol Lett 333, 1–9 (2012).

7. B. Flemer et al., Tumour-associated and non-tumour-associated microbiota in colorectal cancer. Gut 66, 633–643 (2017).

8. R. V. Purcell, M. Visnovska, P. J. Biggs, S. Schmeier, F. A. Frizelle, Distinct gut microbiome patterns associate with consensus molecular subtypes of colorectal cancer. Sci Rep 7, 11590 (2017).

9. W. Mu et al., Intracellular Porphyromonas gingivalis Promotes the Proliferation of Colorectal Cancer Cells via the MAPK/ERK Signaling Pathway. Front Cell Infect Microbiol 10, 584798 (2020).

10. S. Okumura et al., Gut bacteria identified in colorectal cancer patients promote tumourigenesis via butyrate secretion. Nat Commun 12, 5674 (2021).

11. X. Wang et al., Porphyromonas gingivalis Promotes Colorectal Carcinoma by Activating the Hematopoietic NLRP3 Inflammasome. Cancer Res 81, 2745–2759 (2021).

12. G. Lattanzi et al., iNKT cell-neutrophil crosstalk promotes colorectal cancer pathogenesis. Mucosal Immunol 16, 326–340 (2023).

13. C. M. Crosby, M. Kronenberg, Tissue-specific functions of invariant natural killer T cells. Nat Rev Immunol 18, 559–574 (2018).

14. G. Delfanti, P. Dellabona, G. Casorati, M. Fedeli, Adoptive Immunotherapy With Engineered iNKT Cells to Target Cancer Cells and the Suppressive Microenvironment. Front Med (Lausanne) 9, 897750 (2022).

15. Z. Metzger et al., Synergistic pathogenicity of Porphyromonas gingivalis and Fusobacterium nucleatum in the mouse subcutaneous chamber model. J Endod 35, 86–94 (2009).

16. E. Germanov et al., Critical role for the chemokine receptor CXCR6 in homeostasis and activation of CD1d-restricted NKT cells. J Immunol 181, 81–91 (2008).

17. M. Gershkovitz et al., TRPM2 Mediates Neutrophil Killing of Disseminated Tumor Cells. Cancer Res 78, 2680–2690 (2018).

18. C. Burrello et al., Mucosa-associated microbiota drives pathogenic functions in IBD-derived intestinal iNKT cells. Life Sci Alliance 2, (2019).

19. A. Diaz-Basabe et al., Human intestinal and circulating invariant natural killer T cells are cytotoxic against colorectal cancer cells via the perforin-granzyme pathway. Mol Oncol 15, 3385–3403 (2021).

20. T. Zhao, Z. Su, Y. Li, X. Zhang, Q. You, Chitinase-3 like-protein-1 function and its role in diseases. Signal Transduct Target Ther 5, 201 (2020).

21. A. Darwich et al., Paralysis of the cytotoxic granule machinery is a new cancer immune evasion mechanism mediated by chitinase 3-like-1. J Immunother Cancer 9, (2021).

22. Q. Sun et al., STAT3 regulates CD8+ T cell differentiation and functions in cancer and acute infection. J Exp Med 220, (2023).

23. C. Burrello et al., Fecal Microbiota Transplantation Controls Murine Chronic Intestinal Inflammation by Modulating Immune Cell Functions and Gut Microbiota Composition. Cells 8, (2019).

24. C. Burrello et al., IL10 Secretion Endows Intestinal Human iNKT Cells with Regulatory Functions Towards Pathogenic T Lymphocytes. J Crohns Colitis 16, 1461–1474 (2022).

25. F. Strati et al., Antibiotic-associated dysbiosis affects the ability of the gut microbiota to control intestinal inflammation upon fecal microbiota transplantation in experimental colitis models. Microbiome 9, 1–15 (2021).

26. T. Tachibana et al., Increased intratumor Valpha24-positive natural killer T cells: a prognostic factor for primary colorectal carcinomas. Clin Cancer Res 11, 7322–7327 (2005).

27. L. S. Metelitsa et al., Natural killer T cells infiltrate neuroblastomas expressing the chemokine CCL2. J Exp Med 199, 1213–1221 (2004).

28. D. E. Gaddis, C. L. Maynard, C. T. Weaver, S. M. Michalek, J. Katz, Role of TLR2-dependent IL-10 production in the inhibition of the initial IFN-gamma T cell response to Porphyromonas gingivalis. J Leukoc Biol 93, 21–31 (2013).

29. N. M. Moutsopoulos et al., Porphyromonas gingivalis promotes Th17 inducing pathways in chronic periodontitis. J Autoimmun 39, 294–303 (2012).

30. Y. Cai, R. Kobayashi, T. Hashizume-Takizawa, T. Kurita-Ochiai, Porphyromonas gingivalis infection enhances Th17 responses for development of atherosclerosis. Arch Oral Biol 59, 1183–1191 (2014).

31. I. Glowczyk et al., Inactive Gingipains from P. gingivalis Selectively Skews T Cells toward a Th17 Phenotype in an IL-6 Dependent Manner. Front Cell Infect Microbiol 7, 140 (2017).

32. C. Zenobia, G. Hajishengallis, Porphyromonas gingivalis virulence factors involved in subversion of leukocytes and microbial dysbiosis. Virulence 6, 236–243 (2015).

33. D. H. Kim et al., Regulation of chitinase-3-like-1 in T cell elicits Th1 and cytotoxic responses to inhibit lung metastasis. Nat Commun 9, 503 (2018).

34. B. Geng et al., Chitinase 3-like 1-CD44 interaction promotes metastasis and epithelial-to-mesenchymal transition through beta-catenin/Erk/Akt signaling in gastric cancer. J Exp Clin Cancer Res 37, 208 (2018).

35. C. Guetta-Terrier et al., Chi3l1 Is a Modulator of Glioma Stem Cell States and a Therapeutic Target in Glioblastoma. Cancer Res 83, 1984–1999 (2023).

36. Z. Shan et al., Chitinase 3-like-1 contributes to acetaminophen-induced liver injury by promoting hepatic platelet recruitment. Elife 10, (2021).

37. J. D. Klement et al., An osteopontin/CD44 immune checkpoint controls CD8+ T cell activation and tumor immune evasion. J Clin Invest 128, 5549–5560 (2018).

38. M. Krasilnikov, V. N. Ivanov, J. Dong, Z. Ronai, ERK and PI3K negatively regulate STAT-transcriptional activities in human melanoma cells: implications towards sensitization to apoptosis. Oncogene 22, 4092–4101 (2003).

39. S. Zou et al., Targeting STAT3 in Cancer Immunotherapy. Mol Cancer 19, 145 (2020).

40. F. Caprioli et al., Autocrine regulation of IL-21 production in human T lymphocytes. J Immunol 180, 1800–1807 (2008).

41. J. Cui et al., Requirement for Valpha14 NKT cells in IL-12-mediated rejection of tumors. Science 278, 1623–1626 (1997).

42. C. Becker, M. C. Fantini, M. F. Neurath, High resolution colonoscopy in live mice. Nat Protoc 1, 2900–2904 (2006).

43. J. Brummelman et al., Development, application and computational analysis of high-dimensional fluorescent antibody panels for single-cell flow cytometry. Nat Protoc 14, 1946–1969 (2019).

44. V. Bianchi et al., Integrated Systems for NGS Data Management and Analysis: Open Issues and Available Solutions. Front Genet 7, 75 (2016).

45. M. I. Love, W. Huber, S. Anders, Moderated estimation of fold change and dispersion for RNA-seq data with DESeq2. Genome Biol 15, 550 (2014).

46. B. T. Sherman et al., DAVID: a web server for functional enrichment analysis and functional annotation of gene lists (2021 update). Nucleic Acids Res, (2022).

47. P. Bella et al., Blockade of IGF2R improves muscle regeneration and ameliorates Duchenne muscular dystrophy. EMBO Mol Med 12, e11019 (2020).

